# Hoverfly (*Eristalis tenax*) pursuit of artificial targets

**DOI:** 10.1101/2022.07.27.501787

**Authors:** Malin Thyselius, Yuri Ogawa, Richard Leibbrandt, Trevor J. Wardill, Paloma T. Gonzalez-Bellido, Karin Nordström

## Abstract

The ability to visualize small moving objects is vital for the survival of many animals, as these could represent predators or prey. For example, predatory insects, including dragonflies, robber flies and killer flies, perform elegant, high-speed pursuits of both biological and artificial targets. Many non-predatory insects, including male hoverflies and blowflies, also pursue targets during territorial or courtship interactions. To date, most hoverfly pursuits were studied outdoors. To investigate naturalistic hoverfly (*Eristalis tenax*) pursuits under more controlled settings, we constructed an indoor arena that was large enough to encourage naturalistic behavior. We presented artificial beads of different sizes, moving at different speeds, and filmed pursuits with two cameras, allowing subsequent 3D reconstruction of the hoverfly and bead position as a function of time. We show that male *E. tenax* hoverflies are unlikely to use strict heuristic rules based on angular size or speed to determine when to start pursuit, at least in our indoor setting. We found that hoverflies pursued faster beads when the trajectory involved flying downwards towards the bead. Furthermore, we show that target pursuit behavior can be broken down into two stages. In the first stage the hoverfly attempts to rapidly decreases the distance to the target by intercepting it at high speed. During the second stage the hoverfly’s forward speed is correlated with the speed of the bead, so that the hoverfly remains close, but without catching it. This may be similar to dragonfly shadowing behavior, previously coined ‘motion camouflage’.

## Introduction

The ability to visually detect small moving objects can be essential for survival, as these could represent predators or prey. Such visual identification of prey is common in vertebrates, including zebrafish larvae (Mearns et al., 2020; Patterson et al., 2013), archerfish (Newport and Schuster, 2020) and birds of prey (Kane et al., 2015). Visually driven, high-performance predatory attacks are also displayed by insects, including robber flies (Wardill et al., 2017), dragonflies (Olberg et al., 2000), killer flies (Wardill et al., 2015) and praying mantises (Nityananda et al., 2018). Killer flies (Wardill et al., 2015), some robber flies (Fabian et al., 2018) and dragonflies (e.g. Olberg et al., 2005; Olberg et al., 2000), detect their prey from a perch before launching a high-speed pursuit to catch their prey in mid-air. Killer flies decide whether to attack based on the ratio between the angular size of the target image and its angular speed (Wardill et al., 2015). Libelulid dragonflies also use heuristic cues based on the target’s angular size and speed (Lin and Leonardo, 2017).

Non-predatory *Eristalis sp*. hoverfly males pursue both conspecifics and heterospecifics encountered within their territories (Wellington and Fitzpatrick, 1981), but also other small moving targets presented to them (Fitzpatrick, 1981). As opposed to pursuits in more widely studied dragonflies (Olberg et al., 2000), killer flies (Wardill et al., 2015) and robber flies (Wardill et al., 2017), *Eristalis sp*. hoverflies often start their pursuit from a hovering or flying stance (Collett and Land, 1975b; Fitzpatrick, 1981; Wellington and Fitzpatrick, 1981). It has been suggested that hoverflies also use heuristic rules to determine when to initiate pursuit (Collett and Land, 1978). Indeed, based on conspecific size and typical flight speed (Golding et al., 2001; Nationalnyckeln, 2009), the male hoverfly can predict the expected angular size and speed at suitable distances (Collett and Land, 1978). In field observations *Eristalis sp*. males pursue artificial targets with a similar size to conspecifics (Collett and Land, 1978; Fitzpatrick, 1981; Fitzpatrick and Wellington, 1983; Maier and Waldbauer, 1979; Wellington and Fitzpatrick, 1981) moving at 5 – 12.5 m/s (Collett and Land, 1978). However, female *Eristalis sp*. often move much slower than this (Thyselius et al., 2018), indicating that there are instances when such heuristic rules may be broken.

Interestingly, *Eristalis sp*. hoverflies also pursue objects that are much larger than conspecifics, such as leaves (Maier and Waldbauer, 1979), butterflies, hornets and bumblebees (Wellington and Fitzpatrick, 1981), arguing against the use of strict heuristic rules based on angular size. Other fly species also pursue targets that do not appear to be ecologically relevant. Predatory killer flies and non-predatory blowflies may pursue targets that are 3 – 5 times their own size (Boeddeker et al., 2003; Lyneborg et al., 1975; Solano-Rojas et al., 2017; Wardill et al., 2015). However, when blowflies pursue larger beads, they fly further away, consistent with angular size based heuristic rules. If hoverflies similarly adjust their flight behavior to the target’s angular size, is currently unknown.

Following pursuit start, killer flies and robber flies intercept the target using proportional navigation by keeping the bearing angle constant (Fabian et al., 2018). In contrast, blowflies and houseflies use smooth pursuit, by correlating their yaw rotation with the target error angle (Boeddeker et al., 2003; Land and Collett, 1974; Wehrhahn et al., 1982). More recent studies show that blowflies have two different pursuit strategies (Varennes et al., 2020). In the horizontal plane they fly towards the current position of the target, i.e. they use smooth pursuit by aiming to keep the error angle close to anterior, whereas in the vertical plane they use proportional navigation (Varennes et al., 2020). Some hoverflies, like *Syritta*, also use smooth pursuit based on the target error angle (Collett and Land, 1975a), whereas the larger *Eristalis* and *Volucella* intercept the target using deviated pursuit, where the pursuer flies towards the predicted future position of the target by keeping the error angle constant (Collett and Land, 1978).

To investigate the nuances of hoverfly target responses with higher resolution and behavioral control, we developed an indoor arena to record conspecific flight chases in the lab. By making it large enough to encourage naturalistic behavior, we could reconstruct pursuits of beads of different sizes (6 mm to 38.5 mm diameter). We found that *E. tenax* males pursued these artificial targets in the arena, moving at speeds up to 2 m/s, starting their pursuit from the wing either below or above the bead. We found that pursuits were initiated across a large range of angular sizes and speeds, arguing against the use of strict heuristic rules. We show that the hoverflies first accelerate to quickly get close to the artificial target, and then follow the target at a closer distance for up to several seconds. When the hoverfly is proximal to the target its translational speed is correlated with the bead speed, but not with its angular size or speed. Furthermore, we found that flight behavior was different when pursuing the 38.5 mm diameter bead, compared to the beads that were more similar in size to conspecifics. Indeed, hoverflies initiate pursuit of the 38.5 mm bead when it moved slower, they spent longer time distal to it, and they interacted with it physically more often.

## Materials and methods

### Animals

*Eristalis tenax* (Syrphidae) hoverflies were reared from farm collected larvae as described previously (Nicholas et al., 2018). At any one time, we kept ten male and eight female *E. tenax* hoverflies in the flight arena (see below), with females present to encourage male competition. Males showed an interest in females about two weeks after emerging from the pupae and started pursuing artificial targets about two months after emerging.

Hoverflies had constant access to fresh pollen-sugar mix and water. A total of 94 males and 80 female hoverflies lived in the arena until death, or until the time when their physical activity declined noticeably, when they were replaced, to keep the total number constant. Replacement hoverflies came from aged matched, or slightly younger, artificial hibernation stock (Nicholas et al., 2018). In addition, all hoverflies were changed four times (Table S1).

### Flight arena and videography

We used a Plexiglass arena (1 m^3^, Fig. 1A, Movie S1-3), custom designed by Akriform Plast AB (Sollentuna, Sweden). The arena was lit from above by two daylight fluorescent lamps (58 W/865, Nova Group AB, Helsingborg, Sweden) and two office fluorescent lamps, giving an average illuminance of 900 – 1200 lux (LM-120 Light Meter, Amprobe, Everet, WA, USA). The hoverflies were kept at a 12:12 light:dark cycle at room temperature (19 – 20°C).

**Figure 1.**
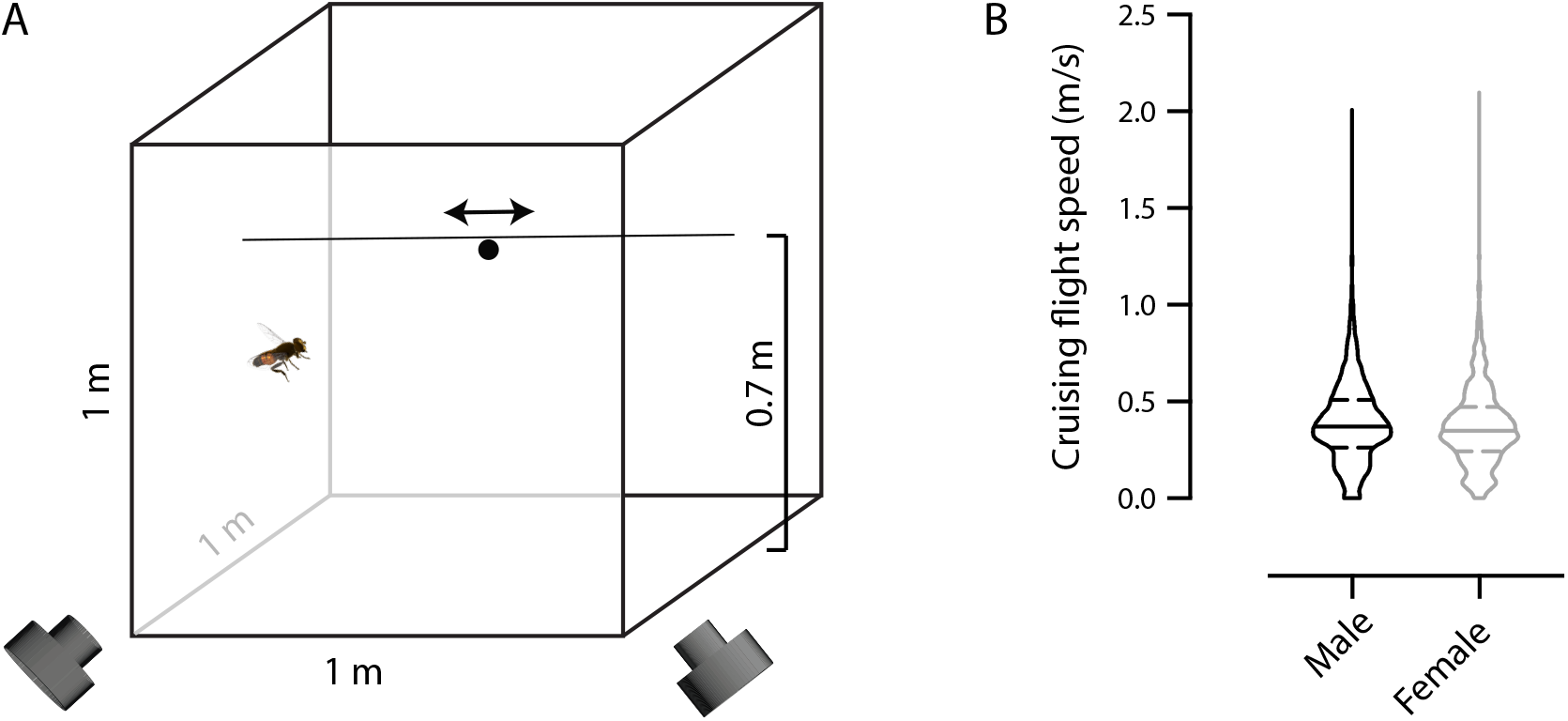
An indoor arena. A) Schematic diagram of the flight arena consisting of a Plexiglas cube (1 m^3^) with a fishing line that formed a horizontal path 0.7 m above the arena floor, with bead movement controlled by a rotor (not shown). For clarity, the hoverfly and bead are shown out of proportion and only one of the 18 hoverflies shown. Experiments were filmed at 120 frames/s using two synchronized cameras (front). B) Hoverfly flight speed at each time point during 100 consecutive frames (0.83 s) of cruising for 10 male (black) and 10 female (grey) flights. Horizontal solid lines show the median and the dashed lines show the interquartile ranges.

A fishing line (0.3 mm diameter) was looped around the arena, entering horizontally through two holes at 0.7 m height (Fig. 1A, Movie S1-3), with a bead attached using a 0.06 mm diameter fishing line. A laptop controlled a stepper motor via a stepper driver (23HS-108 MK.2 Stepper motor and ST5-Q-NN DC input stepper driver, Promoco Scandinavia AB, Täby, Sweden), similar to previously (Wardill et al., 2017). We used ten programs with seven speeds each (0.1 - 2 m/s) presented in a randomized order. Which program was used on any given day was randomized using a 10-sided die. The bead started its motion 10 cm from one side of the arena (Fig. 1A), travelled 0.8 m to the other side, paused for 0.5 s, travelled back to its start, paused for 0.5 s, and then repeated the motion at a different speed (Movie S1-3). The program was run continuously, pausing 3 s before looping.

We used beads of four sizes (6, 8, 10 and 38.5 mm in diameter), painted glossy black (acrylic paint). The three smaller beads were made of glass (Panduro Hobby AB, Malmö, Sweden) and the 38.5 mm diameter bead of polystyrene (Clas Ohlson, Insjön, Sweden). The 6 mm and 10 mm diameter beads were used for 9 consecutive days each. The 38.5 mm bead was used in two periods of 7 and 2 consecutive days. The 8 mm bead was used for 10 consecutive days, and in addition as a control before and after the 6 mm bead experiments, and between the two periods of using the 38.5 mm bead.

We filmed for 39 min ± 2 min (mean ± SEM) per day using two cameras (120 frames/s, 640×480 pixels; EXFH25, Casio, Tokyo, Japan). Filming was done continuously, pausing briefly every 10 minutes to start a new movie, due to saving constraints. The cameras were placed about 1.5 m from the front of the arena, with a 2.5 m distance between the two. They were placed on tripods (Dörr cybrit medi 4-BA, Dörr GmbH, Neu-Ulm, Germany; SIRUI T-2005X, SIRUI, Verona, NJ, USA) with each camera facing the arena at an angle of about 45°, with a 90° angle between the two. The cameras had a focal length of 26 mm, with the focus adjusted to the center of the arena, level with the bead track. In the resulting movies, in the center of the arena 1 pixel corresponds to 3 mm in both the x- and y-plane. As *Eristalis* hoverflies have an average width of about 4 mm and a length of 1.4 cm (Nationalnyckeln, 2009), when far away from the camera, they would sometimes only cover one pixel in an individual frame.

The cameras were synchronized to a 1-frame resolution using the flashlight of a Samsung Galaxy A3 2017 A320 mobile phone (Thyselius et al., 2018; Wardill et al., 2017). The cameras were calibrated as described previously (Thyselius et al., 2018; Wardill et al., 2017) using a 35 mm checkerboard pattern, printed on a polyester sheet and glued to a board (NeverTear, Arkitektkopia AB, Stockholm, Sweden).

### Pursuit and cruising flights

We first identified cruising flights as those where a hoverfly was flying around but clearly not interacting with the bead, another hoverfly, or the arena walls (similar to Collett and Land, 1975a). We arbitrarily selected ten cruising flights from male hoverflies, and ten from females, each 0.83 s (100 frames) long. We next manually identified video recordings where a male hoverfly clearly flew towards the bead and labelled these as visually identified pursuits (Table 1, Movie S1-3).

**Table 1.**
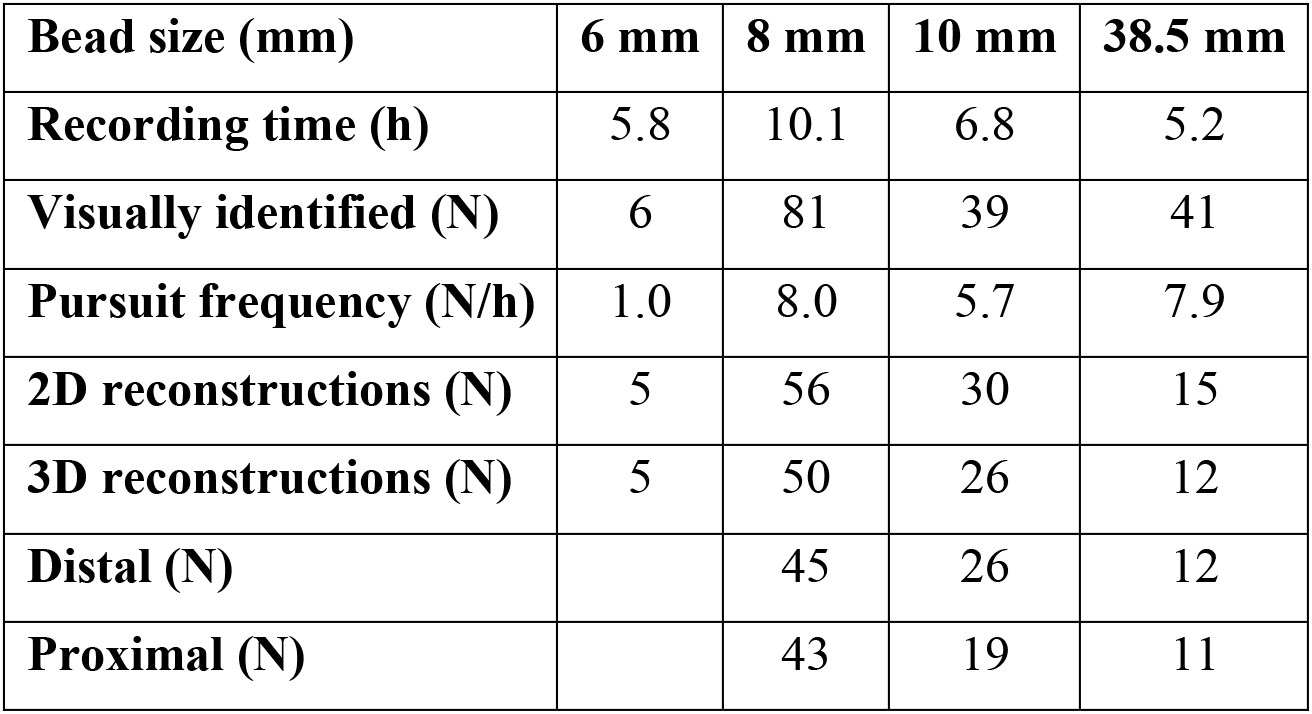
Number of pursuits at each stage of the analysis.

| **Bead size (mm)** | **6 mm** | **8 mm** | **10 mm** | **38.5 mm** |
| --- | --- | --- | --- | --- |
| **Recording time (h)** | 5.8 | 10.1 | 6.8 | 5.2 |
| **Visually identified (N)** | 6 | 81 | 39 | 41 |
| **Pursuit frequency (N/h)** | 1.0 | 8.0 | 5.7 | 7.9 |
| **2D reconstructions (N)** | 5 | 56 | 30 | 15 |
| **3D reconstructions (N)** | 5 | 50 | 26 | 12 |
| **Distal (N)** |  | 45 | 26 | 12 |
| **Proximal (N)** |  | 43 | 19 | 11 |

From each camera frame, we tracked the 2D position of the hoverfly and the bead centroids using custom written Matlab scripts (Thyselius et al., 2018; Wardill et al., 2017), modified to facilitate visual control and manual corrections. We first established the 2D position of the hoverfly and the bead in each frame for each camera (as in Thyselius et al., 2018). Due to the limited spatial resolution only one point was tracked on each. We performed 2D reconstructions (Table 1) of all visually identified pursuits where we could clearly see the hoverfly and the bead throughout. Pursuits where several hoverflies were interacting with each other and/or the bead were excluded.

The 2D data was smoothed using Matlab’s lowess smoothing, which is a locally weighted linear regression method, using 2% of the total number of data points. We translated the 2D positions to 3D (Table 1, Fig. 2A, B), using calibration files obtained from filming the checkerboard, synchronized with both cameras, and previously described methods (Thyselius et al., 2018; Wardill et al., 2017). The 3D data were smoothed again, using 2% of the data points. At this stage, eight pursuits were excluded due to calibration issues, and five because the start of the trajectory was not included, the hoverfly hit the arena wall, or several hoverflies were interacting.

**Figure 2.**
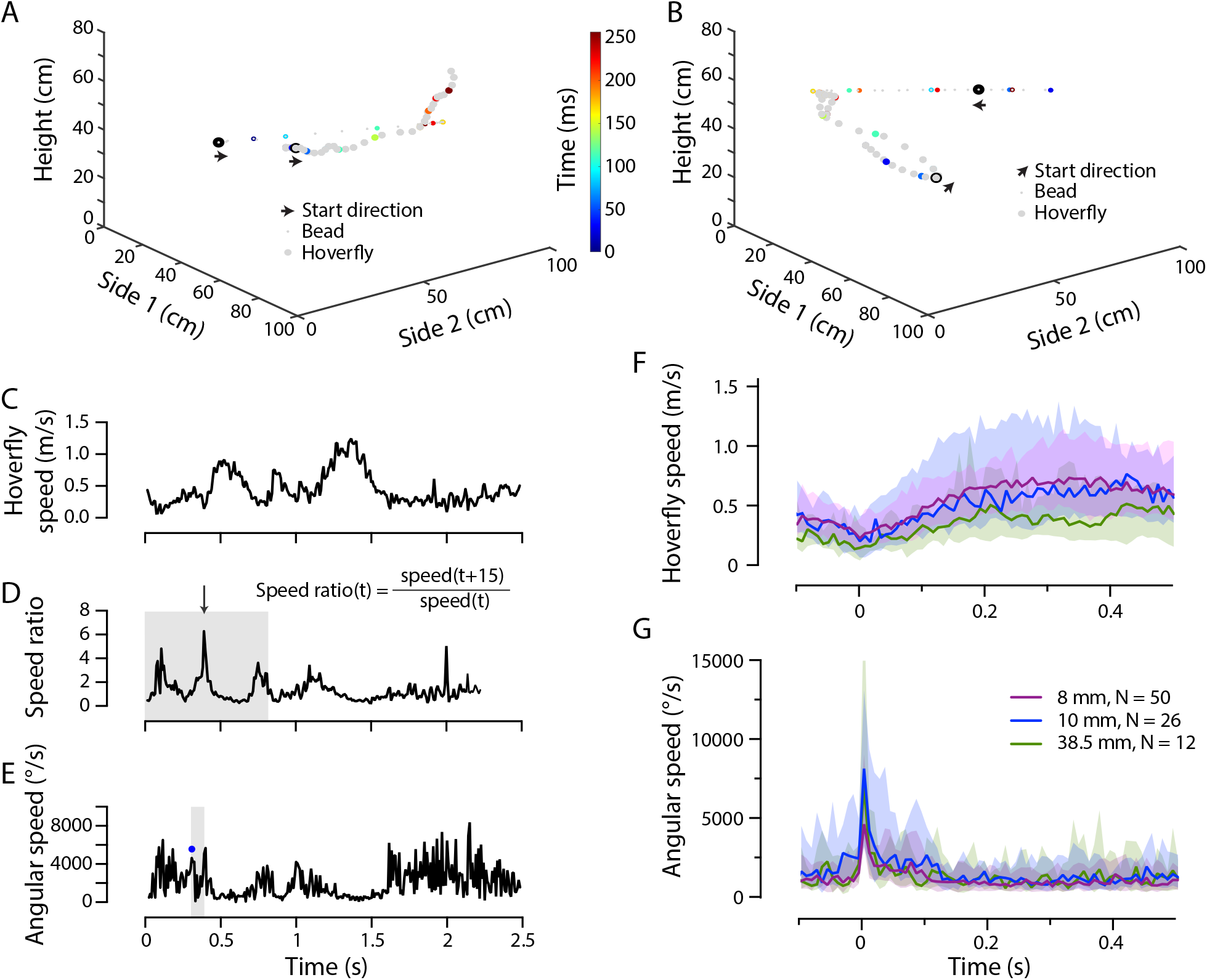
Pursuit start definition. A) An example of a male *E. tenax* (larger symbols) pursuing an 8 mm diameter bead (smaller symbols), displayed at 75 ms resolution. The location of the bead and the hoverfly is color coded every 250 ms. Pursuit start is highlighted with arrows in the direction of travel. B) A second pursuit example. C) The hoverfly’s translational speed of the pursuit example in panel A, as a function of time. D) The speed ratio of the same data (since the speed ratio is calculated across 15 frames, this is 125 ms shorter). We identified the largest peak (arrow) in the first third of the pursuit (shaded). E) The hoverfly’s angular speed during the same pursuit. We defined pursuit start as the largest and sharpest peak (blue symbol) in the 83 ms preceding the speed ratio peak (shaded). F) The hoverflies’ translational speed during pursuit. The data show median +/− interquartile range, color coded according to bead size, with no significant effect of bead size (mixed-effects model). G) The hoverflies’ angular speed during pursuit. The data show the median +/− interquartile range, with no significant effect of bead size (mixed-effects model).

### Data quantification

In all equations below the x- and y-axes define the two sides, and the z-axis the height (Fig. 1A, 2A, B). *F* is used to describe the hoverfly and *B* the bead. The *frame rate* is 120 frames/s, and *t* subsequently defines time steps of 8.3 ms.

As in previous accounts (Collett and Land, 1975b; Varennes et al., 2019), we identified pursuit start based on a sharp turn followed by an increase in translational speed. For this, we first quantified the hoverfly translational speed (*F_s_*), using the Euclidian distance formula, between two consecutive frames:

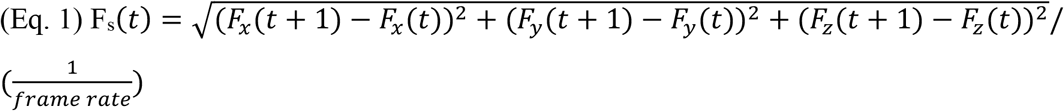

From the hoverfly translational speed (Eq. 1, Fig. 2C) we quantified the speed ratio over 15 frames (125 ms, Fig. 2D):

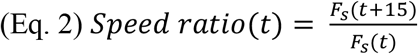

We identified the largest speed ratio peak in the first third of the reconstructed pursuit (grey shading, Fig. 2D). When the bead was stationary or the reconstructed pursuit was longer than 500 frames (4.2 s), we identified the largest speed ratio peak within the first 180 frames (1.5 s).

To determine the hoverfly angular speed, we first calculated its flight heading based on its location change:

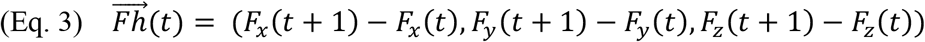

We used the heading change to quantify the hoverfly’s angular speed:

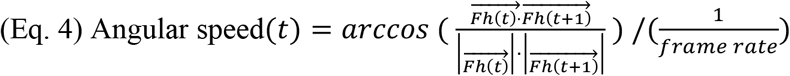

We identified the largest angular speed peak in the 10 frames (83 ms) immediately preceding the speed ratio peak identified above (Eq. 2, grey shading, Fig. 2E). The time of the angular speed peak was used as pursuit start (i.e. *t = 0*).

We smoothed the hoverfly speed (*Fs*, Eq. 1) using a moving average with a span of 10% of the data points, before calculating acceleration:

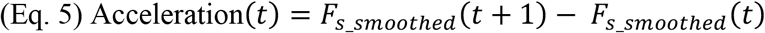

The distance (*d*) was calculated using the formula for Euclidian distance and the 3D coordinates for the hoverfly (*F*) and bead (*B*):

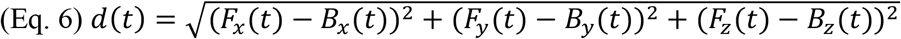

The angular size of the bead image (θ) relative to the hoverfly’s location, was calculated as:

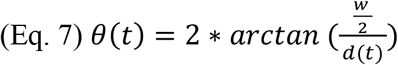

Where *w* was the bead’s physical diameter (8 mm, 10 mm or 38.5 mm) and *d*(*t*), the distance between the bead and hoverfly (Eq. 6).

We quantified the bead speed (*B_s_*) from the bead’s (*B*) 3D position:

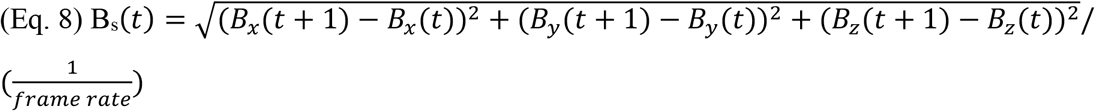

We calculated the relative speed between the hoverfly and the bead using the following:

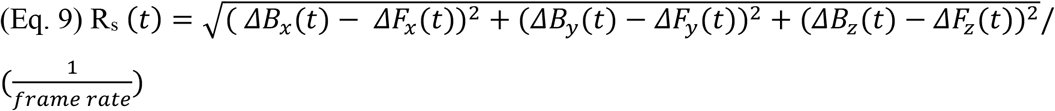

Where *ΔB*_x_(*t*) etc refers to the change in position from the previous frame.

The angular speed of the bead (φ) relative to the hoverfly’s position, was calculated using the law of cosine:

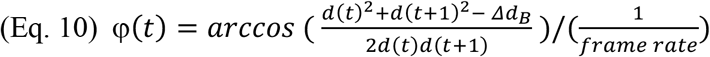

Where *Δd_B_* was the distance the bead travelled relative to the hoverfly’s position from time *t* to time *t+1*. This data was smoothed using a span of 10% of the total number of data points.

We used the distance between the hoverfly and the bead to separate each pursuit into ‘Distal’ and ‘Proximal’. We first determined the distribution of distances across all pursuits, at each time point from pursuit start, for each bead size. We used the lower quartile as a cut-off for the proximal part of the pursuit, i.e. 13.7 cm for the 8 mm bead, 11.9 cm for the 10 mm bead and 8.9 cm for the 38.5 mm bead (Fig. 5D). If the hoverfly was further away than this cut-off at pursuit start, distal was defined as the time between pursuit start and the time when the hoverfly was last outside this cut-off. If the hoverfly was never within the cut-off distance, the minimum distance defined the end of the distal stage. Proximal was identified as the time from when the hoverfly was first within the cut-off distance, until it left it for more than 200 ms.

When calculating the minimum distance between the bead and the hoverfly (Fig. 6), we subtracted the bead radius from *d_minimum_* (Eq. 6).

We defined error angles separately in azimuth and elevation. The error angle (ε) was defined as the 2D angle between the hoverfly heading (*Fh*, Eq. 11a, b) and the Line-of-Sight (*LoS*, Eq. 12a, b). The hoverfly heading vector (*Fh*, Eq. 11a, b) was calculated by subtracting its 2D position between two consecutive frames:

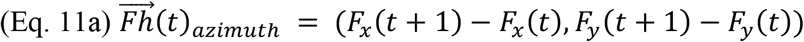

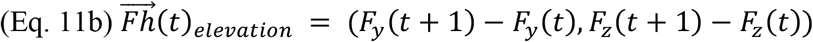

The Line-of-Sight (*LoS*, Eq. 11a, b) vector was defined as the bead’s position relative to the hoverfly’s position:

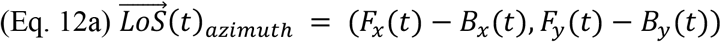

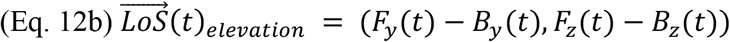

We calculated the 3D error angle as the angle between the hoverfly heading (*Fh*, Eq. 3) and the line of sight (*LoS*, Eq. 13).

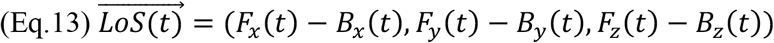

We calculated the 3D bearing angle as the angle between an external reference point (0, 0, −1) and the line of sight (*LoS*, Eq. 13).

All error and bearing angles were smoothed using a span of 4% of the total number of data points. The delta error and delta bearing angles were defined as the absolute change between *t-2* and *t*.

### Visualization and statistics

Data analysis, statistics and figure preparation were done using Prism 9 (GraphPad Software Inc., San Diego, CA, USA) and Matlab (R2019b, The MathWorks, Inc., Natick, MA, USA). All data are shown as either individual data points or median +/− interquartile range, unless otherwise specified. We used a restricted maximum likelihood model to fit mixed-effects models to time aligned data. For pursuit start quantifications, we calculated the mean from *t_−2_* to *t_+2_*, where *t* = pursuit start. For before-start quantifications, we calculated the mean from *t_−18_* to *t_−6_*, where *t* = pursuit start. Since most data were not normally distributed, we used Kruskal-Wallis tests followed by Dunn’s multiple comparisons tests, or Mann-Whitney tests. For circular data we used Matlab’s Circular statistics toolbox (Berens, 2009).

We used the first 1 s of pursuit for all bead sizes to determine the correlation coefficient (using Matlab’s *corrcoef*) for different time shifts (Fig. S3A, B), from which we extracted the time of the peak correlation. For visualization we then extracted the data at each time point for each bead size individually, using the peak correlation delay (Fig. 5G, H). The graphs show the linear regression, together with the R^2^ value or Spearman correlation.

## Results

### Male hoverflies pursue artificial targets in an indoor arena

To assert that *E. tenax* hoverflies behaved naturalistically in our indoor arena (Fig. 1A), we analyzed the flight speed during cruising flights (see Methods). We found that the top cruising speed was just over 2 m/s with a median of 0.37 m/s for males and 0.35 m/s for females (Fig. 1B), which is similar to field flight speeds (0.32 m/s, Golding et al., 2001; 0.34 m/s, Thyselius et al., 2018).

We filmed with two cameras (Fig. 1A) and reconstructed the 3D position of hoverflies pursuing artificial targets. We show two example pursuits (Fig. 2A, B, Movie S1, 2), with the bead (8 mm diameter) and male hoverfly locations every 75 ms. Since hoverflies initiate pursuit while on the wing (Collett and Land, 1975b), we defined pursuit start as a sharp increase in angular speed (i.e. a turn), followed by a translational speed increase (Fig. 2C-E, and see Collett and Land, 1975b; Varennes et al., 2019). We found no difference in translational speed (Fig. 2F) nor angular speed (Fig. 2G) between pursuits of beads of different sizes. The hoverfly acceleration increased after pursuit start (Fig. S1A), but was much lower than the 33m/s^2^ measured in the field (Collett and Land, 1978).

### Target image at pursuit start

We found that male hoverflies pursued black beads of all four sizes (6 mm, 8 mm, 10 mm and 38.5 mm in diameter, Table 1). However, despite often flying in the arena, we only visually identified a few pursuits per hour (Table 1). In addition, pursuits of the 6 mm diameter bead were even more rare than pursuits of the other bead sizes (Table 1, Movie S3) and the data were therefore excluded from further analysis. The hoverfly flight speed 100 ms before pursuit ranged from 0.01 – 1.1 m/s (minimum to maximum), with median values of 0.33 m/s when pursuing the 8 mm bead, 0.45 m/s when pursuing the 10 mm bead, and 0.23 m/s when pursuing the 38.5 mm bead (Fig. S1B).

The hoverflies started pursuit from a range of distances (from 5.6 cm to 91 cm, Fig. 3A), suggesting that they used the entire arena. The median distance at pursuit start was 43 cm for the 8 mm bead, 48 cm for the 10 mm bead, and 44 cm for the 38.5 mm bead (Fig. 3A). The coefficient of variation was 46% for the 8 mm bead, and 41% for both the 10 and 38.5 mm beads (Fig. 3A). The angular size of the bead (inset, Fig. 3B) at pursuit start covered a broad range of 0.5 – 22°, with median sizes for the three bead sizes of 1.1°, 1.2° and 5.0° (Fig. 3B). Consistent with its larger physical size, the larger bead had a significantly larger angular size (green data, Fig. 3B). The angular size coefficients of variation were large, 91%, 66% and 79% (Fig. 3B).

**Figure 3.**
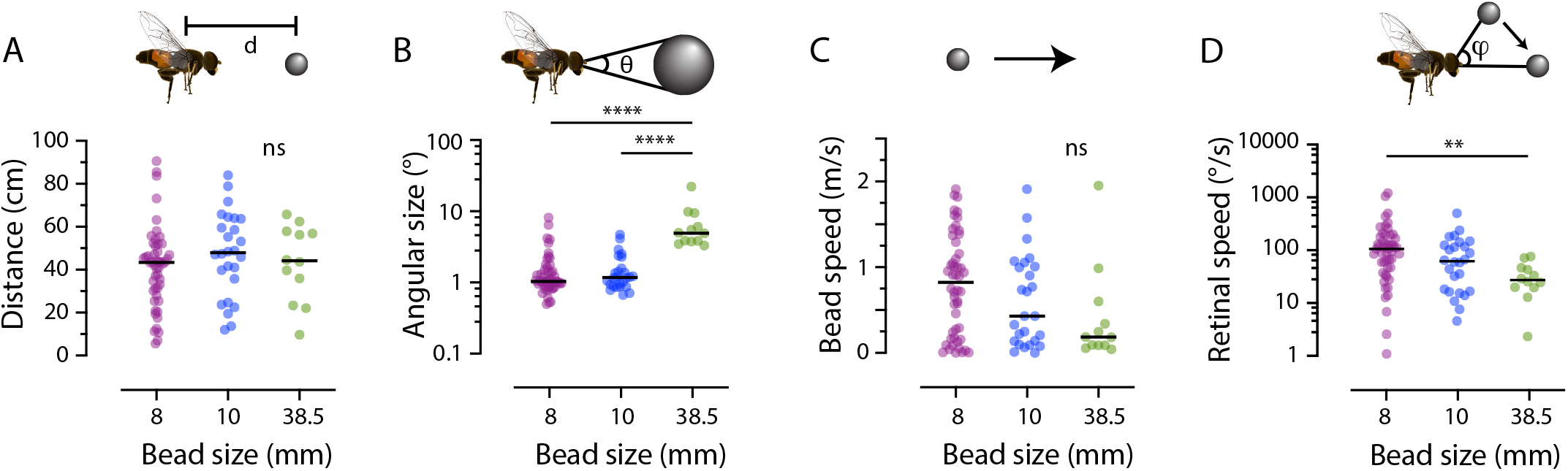
E. tenax *hoverflies do not use strict matched filters to initiate pursuit*. A) The distance between the hoverfly and the bead at pursuit start. B) The angular size of the target image at pursuit start. C) The translational speed of the bead at pursuit start. D) The angular speed of the target image. The data were analyzed using Kruskal-Wallis tests, followed by Dunn’s multiple comparisons, with ** for p < 0.01 and **** for p < 0.0001.

The bead speed at pursuit start ranged from stationary to 2.0 m/s, with coefficients of variation for the three bead sizes of 73%, 86% and 138% (Fig. 3C). The median bead speeds at pursuit start were 0.8 m/s, 0.4 m/s and 0.2 m/s, but this trend was not significant (p = 0.1). We found that the relative speed between the bead and the hoverfly 100 ms before pursuit start ranged from 0.05 to 2.0 m/s, with coefficients of variation of 70%, 64% and 71% (Fig. S1C).

The angular speed of the target image (inset, Fig. 3D) at pursuit start was significantly lower for the largest bead size (Fig. 3D), with median speeds of 94°/s, 79°/s and 36°/s (Fig. 3D). The angular speed coefficients of variation were large, 153%, 97% and 73% (Fig. 3D).

Taken together, due to the large coefficients of variation, it is unlikely that male *E. tenax* hoverflies use strict heuristic rules based on the target’s angular size or speed (Fig. 3B, D) to determine when to initiate pursuit. Killer flies use the ratio between the angular size and speed to determine which targets to pursue (Wardill et al., 2015). However, we found that the coefficient of variation for this ratio was also large, 136%, 146% and 72% (not shown). Neither did the hoverflies seem to selectively pursue targets from a narrow range of distances, physical sizes, nor bead speeds (Fig. 3A, C, Fig. S1C).

### Flight behavior at start

The data above (Fig. 3) show that male *E. tenax* hoverflies are unlikely to use strict heuristic rules based on angular size or speed to trigger pursuit initiation. However, visual information from the bead could be used to adjust initial flight behavior. The pursuit start is associated with a translational speed increase (Collett and Land, 1975b; Varennes et al., 2019), here quantified as a speed ratio peak (example trace shown in Fig. 2D). We found that this ratio did not depend on the bead’s physical diameter (Fig. S2A), nor its angular size (Fig. S2B). Neither did the speed ratio peak depend on the bead’s translational speed (Fig. S2C), nor its angular speed (Fig. S2D).

We next looked at the hoverfly’s increased angular speed (example in Fig. 2E) associated with a sharp turn at pursuit start, and found that it did not depend on the bead size (Fig. S2E), its angular size (Fig. S2F), its translational speed (Fig. S2G), nor angular speed (Fig. S2H). In contrast, field work has shown a correlation between the hoverfly’s angular speed at start and the bead’s angular speed, which has been interpreted as an effort to put the target image in the frontal visual field (Collett and Land, 1978). In our experiment, the hoverflies’ angular speed at pursuit start ranged from 460 to 13,000°/s, with median speeds of 2,960°/s for the 8 mm bead, 4,530°/s for the 10 mm bead and 4,230°/s for the 38.5 mm bead (Fig. S2E), similar to turning speeds measured in the field (Collett and Land, 1978).

We quantified the delay between the hoverfly’s peak angular speed (e.g. blue dot, Fig. 2E) and the peak speed ratio (e.g. arrow in Fig. 2D) and found that this did not depend on the physical size of the bead (Fig. S2I) nor its angular size (Fig. S2J). Neither did the delay depend on the bead’s translational speed (Fig. S2K), nor its angular speed (Fig. S2L). The median delays were 25 ms for the 6 mm bead, 17 ms for the 8 mm bead, and 21 ms for the 38.5 mm bead (Fig. S2I), similar to delays measured in the field (Collett and Land, 1975b). In summary, hoverflies showed similar pursuit start behavior to the field, with a sharp turn followed by a translational speed increase about 20 ms later, suggesting that they behaved naturalistically, but the heuristic rules here investigated could not explain how hoverflies controlled this behavior.

### Pursuits may start from above or below the bead

Killer flies pursue artificial beads from above as well as below (Rossoni et al., 2021). We found that this was also the case for hoverflies, across the bead sizes tested (Fig. 4A). We next investigated if the hoverfly starting position (i.e. above vs below the bead) impacted flight behavior, when pursuing the 8 mm bead, as we had most pursuits of this bead size. We found that the starting position did not affect the hoverfly’s speed ratio (Fig. 4B), its angular speed at start of the pursuit (Fig. 4C), nor the delay between the angular and translational speed peaks (Fig. 4D). In contrast, pursuits that started from below the bead were initiated when the bead was significantly further away from the hoverfly (Fig. 4E). It is likely that this was due to the larger space available below the bead track, which was located 70 cm above the arena floor (Fig. 1A). Consistent with the distance difference (Fig. 4E), the angular size of the target image also differed (Fig. 4F). We found that when the hoverflies started their pursuit from above, they pursued faster targets (Fig. 4G). Together, the shorter distance to the target and the faster target speed resulted in a significant difference in the target’s angular speed when comparing above with below starting positions (Fig. 4H).

**Figure 4.**
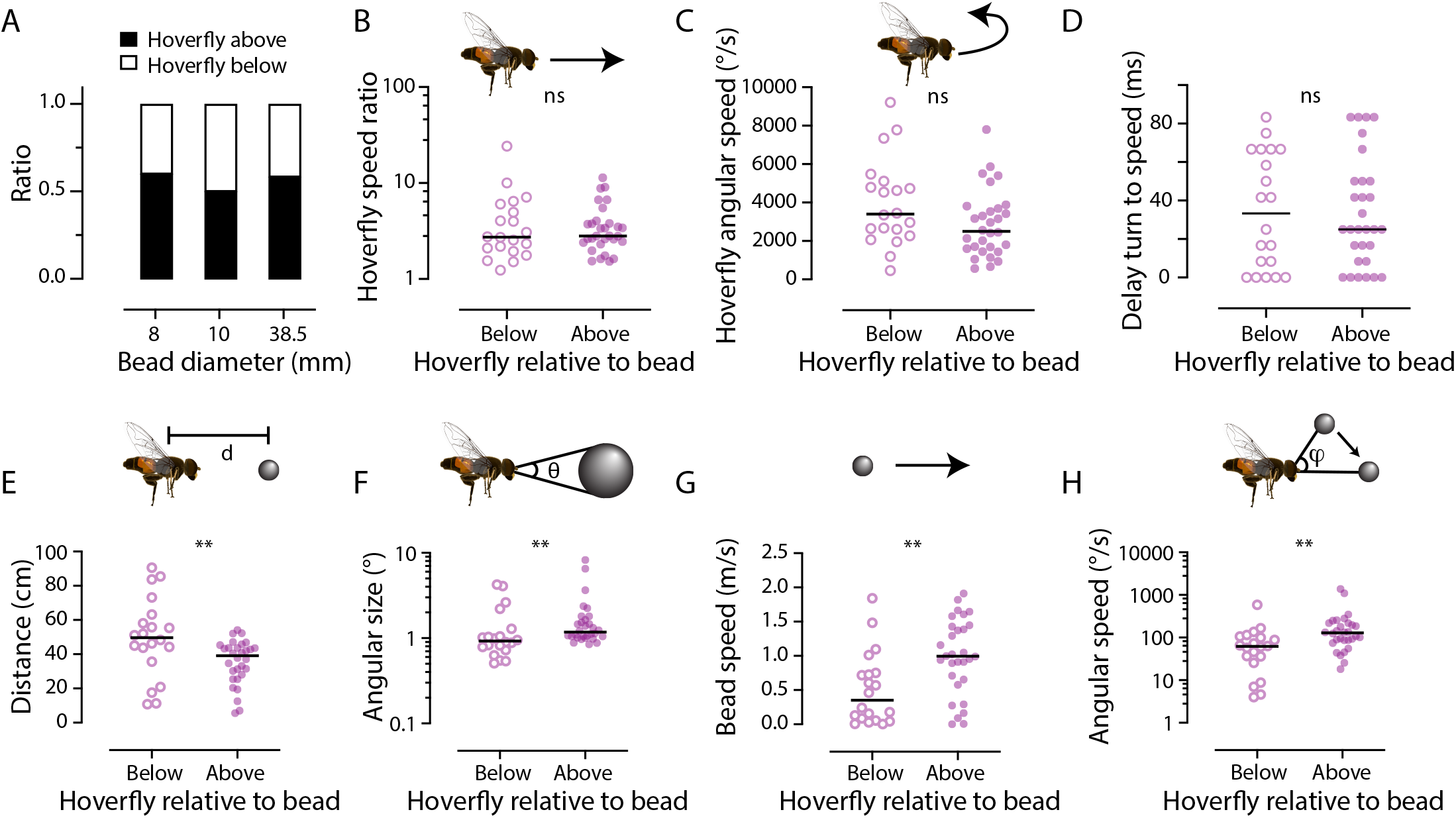
E. tenax *hoverflies initiate pursuits from above as well as below the bead*. A) The elevation of the hoverfly relative to the bead at pursuit start, as a function of bead size (N = 50, 26 and 12 for the three bead sizes). There was no significant difference between the bead sizes (p = 0.93, chi square test). B) The speed ratio of the hoverfly following pursuit start, as a function of whether the hoverfly was below (open symbols) or above (closed symbols) the 8 mm bead. C) The hoverflies’ angular speed at pursuit start, as a function of whether the hoverfly was below or above the bead. D) The delay between the hoverflies’ angular and translational speed ratio peaks. E) The distance between the hoverfly and the bead at pursuit start. F) The angular size of the target image from the hoverfly’s position at pursuit start. G) The translational bead speed at pursuit start. H) The angular speed of the target from the hoverfly’s position at pursuit start. Significance was investigated using Mann-Whitney tests with ** for p < 0.01.

### Male hoverflies divide their pursuits into two stages

We next quantified the hoverfly’s behavior during pursuit. As expected, the distance between the male hoverfly and the bead decreased with time (Fig. 5A), which resulted in the target’s angular size increasing (Fig. 5B). There was no significant effect of bead size on the distance (Fig. 5A), but the angular size was significantly larger for the 38.5 mm diameter bead (Fig. 5B), consistent with its larger physical size. We also measured the angular speed of the target (see inset) and found that there was no significant effect of bead size (Fig. 5C).

**Figure 5.**
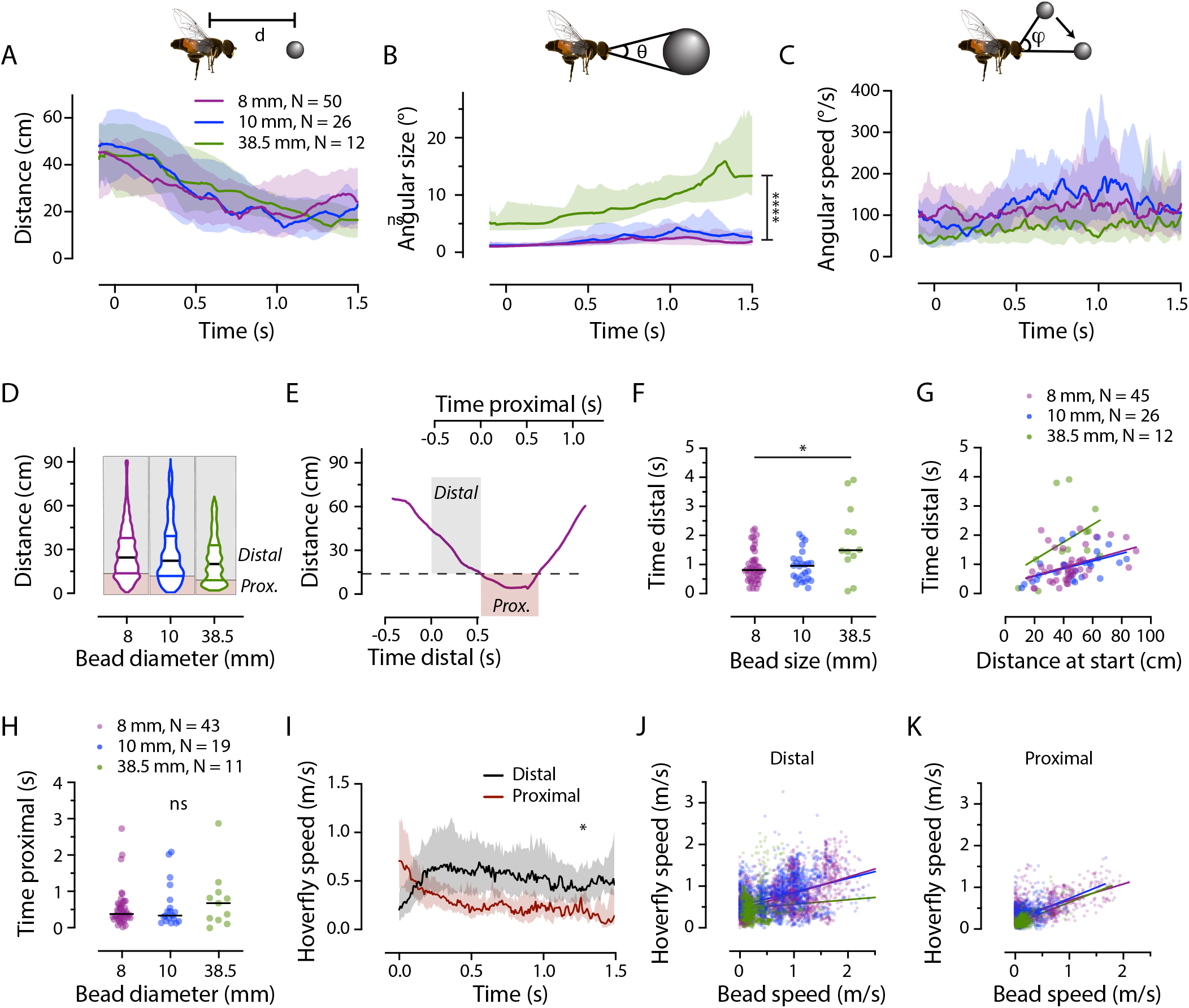
The distal and proximal part of the pursuit. A) The distance between the hoverfly and the bead as a function of time, color coded according to bead size. B) The angular size of the target image. C) The angular speed of the target image. The data in panels A-C were analyzed using mixed-effect models, followed by Tukey’s multiple comparisons test, with **** for p < 0.0001. D) Distances between the hoverfly and the bead at each time point across all pursuits. Horizontal lines show the median (black) and interquartile ranges (colored). The shaded boxes delineate the lower quartile, which was used to separate proximal (brown shading) from distal (grey shading). E) The distance between the hoverfly and the bead for an example pursuit. The time between pursuit start and when last outside the 8 mm bead’s cut-off distance (13.7 cm, dashed line) was defined as distal (grey shading). Proximal (“Prox.”, brown shading) was defined as when within this cut-off distance. F) The time the hoverfly was distal to the bead as function of bead size, analyzed using a Kruskal-Wallis test, followed by Dunn’s multiple comparisons, p = 0.025. G) The time the hoverfly was distal to the bead as function of the distance to the bead at pursuit start, with linear regression lines (R^2^ = 0.17, 0.25 and 0.19). H) The time the hoverfly was proximal to the bead as function of bead size, analyzed using a Kruskal-Wallis test. I) The hoverfly speed as a function of time, color coded to show the distal (black) and proximal part (brown). The data show the median +/− interquartile range, and the * indicates a significant difference (mixed-effect models, followed by Tukey’s multiple comparisons test). J) The hoverflies’ translational speed at each time point as a function of the bead speed 233 ms previously, during the first 1 s of the distal part of the pursuit, with linear regression lines (R^2^ = 0.26, 0.15 and 0.009). K) The hoverflies’ translational speed at each time point as a function of the bead speed 150 ms previously, during the first 1 s of the proximal part of the pursuit, with linear regression lines (R^2^ = 0.35, 0.21 and 0.39).

To investigate the behavior during pursuit further, we first looked into the bead-to-hoverfly distance across all trajectories, at each time point. The bead-to-hoverfly distance range was 0.5 – 92 cm, and the median distances for the three bead sizes were 25, 22 and 20 cm (Fig. 5D). We also noted that across pursuits, more time points were spent close to the bead compared with further away (Fig. 5D). To investigate if the behavior was different when closer to the bead compared with further away, we used the lower quartile for each bead size (13.7, 11.9 and 8.9 cm, Fig. 5D, E) to separate each pursuit into a ‘distal’ (far from bead) and a ‘proximal’ (close to bead) stage (see Methods for details).

We found that the time the hoverfly spent distal to the bead was highly variable, from 83 ms to 3.9 s, with median durations for the three bead sizes of 0.81 s, 0.95 s and 1.5 s, which was significant (Fig. 5F). The total amount of time that the bead-to-hoverfly distance exceeded the distal cutoff, increased linearly with the distance to the bead at pursuit start (R^2^ values of 0.17, 0.25 and 0.19, Fig. 5G). I.e., as may be expected, the further away the fly started from the bead, the longer it spent in the distal phase, but the correlation was not significant for the 38.5 mm bead (p = 0.0024 for the 8 mm bead, p = 0.0032 for the 10 mm bead and p = 0.14 for the 38.5 mm bead, Fig. 5G).

The time the hoverfly was proximal to the bead ranged from 8.3 ms to 2.9 s, with median durations of 0.39 s, 0.35 s and 0.69 s for the three bead sizes, but this difference was not significant (Fig. 5H). We hypothesized that the distal stage was optimized to rapidly decrease the distance to the bead (Fig. 5A, D-G), while the proximal stage (Fig. 5D, E, H) was aimed at staying close to the bead. In support of this, the hoverfly translational speed was higher during the distal stage than during the proximal stage (Fig. 5I).

To investigate how hoverflies control their translational speed during pursuit, we calculated the correlation coefficients between the hoverfly speed and the bead distance, its angular size, the bead speed, and its angular speed, during the first second of the distal stage (Fig. S3A). We found the strongest correlation between the hoverfly’s translational speed and the bead’s speed (solid data, Fig. S3A), with a peak at −233 ms (dotted vertical line, Fig. S3A). The graph showing the flight speed at each time point, as a function of the bead speed 233 ms previously (Fig. 5J), highlights that even if they are correlated, there is large variation, with R^2^ values of 0.26 (purple, 8 mm bead), 0.15 (blue, 10 mm bead), and only 0.010 (green, 38.5 mm bead, Fig. 5J). In addition, 233 ms is slow for typical insect reactions (e.g. Collett and Land, 1978; Mischiati et al., 2015; Varennes et al., 2020; Wehrhahn et al., 1982), so its biological relevance needs to be taken with caution. Together, this suggests that during the distal stage the hoverfly was aiming to rapidly decrease the distance to the bead (Fig. 5A, E, G, I).

We carried out cross correlations for the proximal stage and found the strongest correlation between the hoverfly’s translational speed and the bead’s speed (solid data, Fig. S3B) at −150 ms (dotted vertical line, Fig. S3B). We visualized this by plotting the hoverfly speed as a function of the bead speed 150 ms previously, at each time point, from all the pursuits where the hoverfly was proximal to the bead, and found R^2^ values of 0.35 (purple, 8 mm), 0.21 (blue, 10 mm) and 0.39 (green, 38.5 mm bead, Fig. 5K).

We found a weaker correlation between the hoverfly’s translational speed and the distance to the bead, with a peak at −42 ms (dashed, Fig. S3B). In the graph showing the hoverfly speed as a function of distance 42 ms previously, for each time point, we found R^2^ values of 0.096, 0.11 and 0.006 (Fig. S3C).

To test if the observed reduction in flight speed during the proximal stage (Fig. 5I, K) was due to the hoverfly being close to the arena wall, we plotted the flight speed at each time point as a function of the horizontal distance to the closest arena wall (Fig. S3D). We found that the hoverflies flew fast even close to the walls, making the wall an unlikely confounding factor. Taken together, our findings suggests that in the proximal phase, the hoverfly matched the target speed (Fig. 5K) because it intended to shadow it, rather than catch it.

If true, this should be reflected in the minimum distance between the bead and the hoverfly. Indeed, we found that pursuits of the 8 and 10 mm beads rarely ended with the hoverfly grabbing or landing on the bead (median minimum distances 5.8 and 6.5 cm, Fig. 6). In contrast, the median distance between the hoverfly and the 38.5 mm bead was 0.99 cm (green data, Fig. 6), suggesting that the hoverflies often landed on the largest bead. This is interesting considering that they initiated pursuit of the 38.5 mm bead when it moved slower (Fig. 3C), and they spent longer time distal to it (Fig. 5F). Maybe the hoverflies categorized it differently to the more conspecific-sized targets.

**Figure 6.**
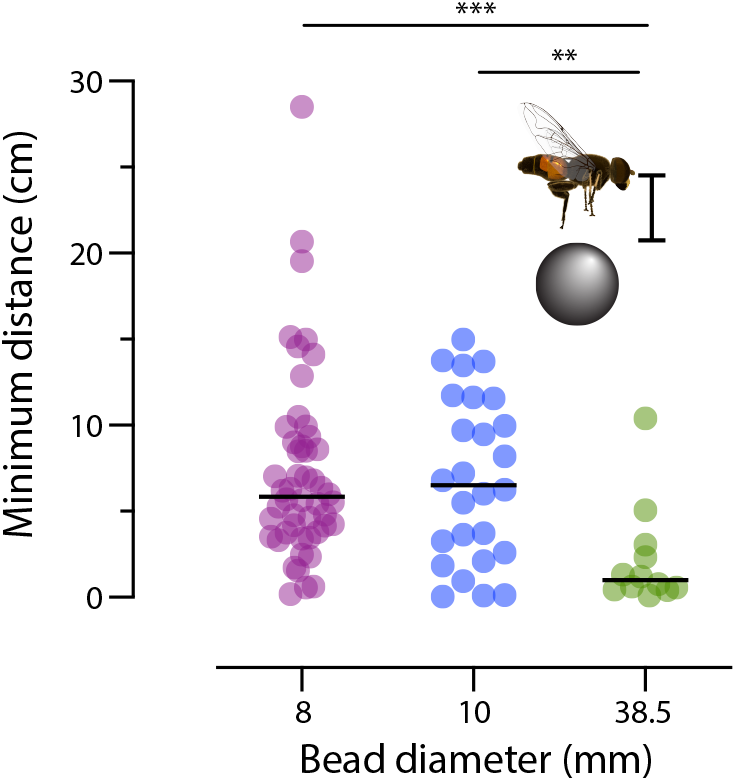
The hoverflies land on the largest bead, but not the others. The minimum distance between the hoverfly and the bead, here defined as the distance between the surface of the bead and the center of the hoverfly. The data were analyzed using Kruskal-Wallis tests, followed by Dunn’s multiple comparisons, with ** indicating p < 0.01 and *** for p < 0.001.

### Error angles during pursuit

As we found that *Eristalis* hoverflies pursue beads from above and below (Fig. 4), next we analyze if the initial geometry between fly and target had an impact on the subsequent pursuit behavior. For this purpose we calculated the error angle (blue, Fig. 7A, B), defined as the angle between the hoverfly heading (black arrows, Fig. 7A, B) and the Line-of-Sight to the bead (dashed line, Fig. 7A, B; see e.g. Land and Collett, 1974; Rossoni et al., 2021; Varennes et al., 2020). The error angle was quantified both in the azimuth (Fig. 7A) and elevation planes (Fig. 7B). After smoothing the data (see Methods) we extracted the error angle at five different time points: 100 ms before pursuit start, at the start of the distal stage, 100 ms into the distal stage, at the start of the proximal stage, and 100 ms into the proximal stage.

**Figure 7.**
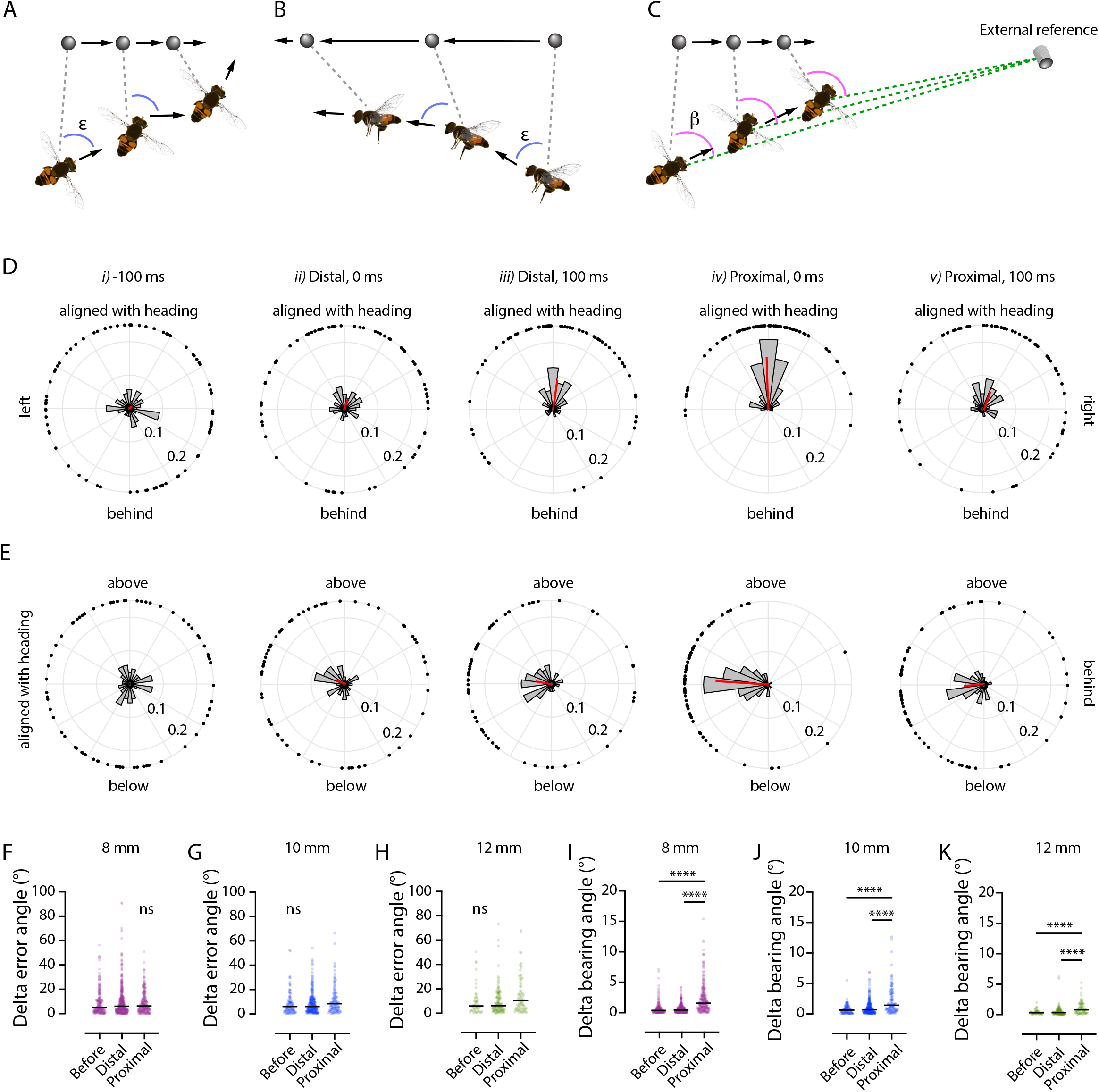
The error and bearing angle. A) We defined the error angle (*ε*, blue) as the angle between the hoverfly’s heading (black arrows) and the direct line connecting the hoverfly’s position and the bead’s location (grey dashed line, often referred to as Line of Sight, LoS). The diagram shows this in the azimuth plane. B) The error angle in the elevation plane. C) The bearing angle (β, magenta) was defined as the angle between the LoS (grey dashed), and an external reference point (green dashed). D) The error angle in the azimuth plane across 5 different time points: *i)* 100 ms before pursuit start (N = 73), *ii)* when distal at pursuit start (N = 83, *iii)* 100 ms into the distal part of the pursuit (N = 82), *iv)* when first being proximal to the bead (N = 73), and *v)* 100 ms into the proximal part of the pursuit (N = 69). E) The elevation error angle across the same 5 time points. In panels C, D the black dots show the error angle for each individual pursuit of the 8, 10 and 38.5 mm diameters beads. The histograms show the distribution of error angles within 15° bins. The red line shows the resulting mean vector. F) The delta error angle, before pursuits of the 8 mm bead, during the distal stage and during the proximal stage. G) The delta error angle, before pursuits of the 10 mm bead, during the distal and the proximal stage. H) The delta error angle, before pursuits of the 38.5 mm bead, during the distal and the proximal stage. I) The delta bearing angle, pursuits of the 8 mm bead. J) The delta bearing angle, pursuits of the 10 mm bead. K) The delta bearing angle, pursuits of the 38.5 mm bead. The data in panels F-K were analyzed using Kruskal-Wallis tests, followed by Dunn’s multiple comparisons, with **** indicating p < 0.0001.

We found that 100 ms before pursuit started, the error angles were evenly distributed, both in the azimuth (Fig. 7D*i*) and the elevation plane (Fig. 7E*i*). This was also the case at the start (0 ms) of the distal stage (azimuth and elevation in Fig. 7D*ii* and 7E*ii*, respectively). This is because hoverflies were flying in different directions when the target caught their attention. We found no correlation between the error angle at pursuit start, and the hoverfly angular speed, for the two smaller bead sizes (blue and purple, Fig. S4), but there was a correlation for the 38.5 mm bead (green data, R^2^ = 0.10, Fig. S4).

We found that 100 ms into the distal stage the mean error angle was directed anteriorly (red, Fig. 7D*iii*, E*iii*). At the start of the proximal stage, the mean error angle was even more strongly anterior (red, Fig. 7D*iii*, E*iv*), but this strong directional preference had decreased after 100 ms (red, Fig. 7D*v*, E*v*). Taken together, our data suggest the hoverflies attempted to adjust their flight direction to keep the target anterior relative to its direction of flight, but there was a large variation (Fig. 7D, E).

Our data above suggested that during the distal stage the hoverflies attempted to rapidly decrease the distance to the targets (Fig. 5E, F, G, I), potentially by intercepting its future position (Fig. 2A, B) as suggested in field work (Collett and Land, 1978). In contrast, during the proximal stage they appeared to follow the speed of the bead more closely (Fig. 5H, I, K), by keeping the target image anterior (Fig. 7D*iv*, E*iv*). Target interception can be achieved by keeping either the error angle (Fig. 7A, B, also referred to as deviated pursuit) or the bearing angle (Fig. 7C, also referred to as proportional navigation) constant. We therefore investigated if the error or bearing angles in 3D space were kept constant, by quantifying how much they changed from one time point to one 2 frames later (16.7 ms). The closer to 0 the delta angle is, the more constant the angle is. We quantified the delta error and delta bearing angles in the 1 s preceding pursuit start, during the distal stage and during the proximal stage.

We found that the delta error angle was not significantly different between the three stages, for any of the bead sizes (Fig. 7F-H). In addition, the delta error angle varied a lot, suggesting that the hoverflies did not attempt to keep it constant, as requested for deviated pursuit (Fig. 7A, B). In contrast, the bearing angle (Fig. 7C), was held much more constant (delta bearing angle close to 0) before the pursuit and during the distal stage, compared with the proximal stage (Fig. 7I-K). This suggests that the hoverflies could use proportional navigation (Fig. 7C) to intercept the bead during the distal stage of the pursuit, whereas they may use smooth pursuit (keeping the error angle close to 0, Fig. 7D*iv*, E*iv*) during the proximal stage. Future modelling endeavors will help elucidate this.

## Discussion

We here showed that *E. tenax* males pursue artificial targets ranging from 6 mm to 38.5 mm in diameter (Table 1) in an indoor flight arena (Fig. 1A, 2, Movie S1-3). We show that male *E. tenax* pursue targets from above as well as below (Fig. 4, 7), with pursuits lasting several seconds (Fig. 5). At the start of the pursuit the hoverflies fly fast to decrease the distance to the bead, whereas they adjust their translational speed to the bead speed when they are proximal (Fig. 5, S3), but only rarely physically interacting with it (Fig. 6). We found that male *E. tenax* are unlikely to use strict heuristic rules based on angular size or speed (Fig. 3, Fig. S2-S4), and that pursuits of the largest bead (green data, Fig. 3D, 5F, 6) differed, suggesting possible categorization.

### Indoor pursuits

The pursuit flight speed in our indoor arena (Fig. 2F, 5I) was lower than the 10 m/s recorded in the field, and the acceleration (Fig. S1A) was also lower than the 33 m/s^2^ measured in the field (Collett and Land, 1978). Therefore, while hoverflies pursued targets in the arena, they were not flying as fast as they do in the field. However, the high angular speeds associated with pursuit start (Fig. 2E, G, S2E) were similar to field measurements (Collett and Land, 1978), suggesting that turning behaviors were naturalistic.

We found it unlikely that hoverflies use strict matched filters, also referred to as heuristic rules, to trigger pursuit start, as the angular size and speed covered a large range of values (Fig. 3B, D). Neither did they seem to adjust their saccade like turn followed by a translational speed increase (Fig. 2C-G, Movie S1-3) at pursuit start to the angular size or speed of the target (Fig. S2, S4), as previously suggested (Collett and Land, 1978). It is possible that being indoors affected the territoriality, and thus reduced the saliency of cues that might be important in the field. Indeed, the pursuit ratio was relatively low (Table 1) compared to field behavior (e.g. Wellington and Fitzpatrick, 1981). Furthermore, having many hoverflies in the arena simultaneously might have added competition, which could affect underlying heuristic rules. Indeed, *Drosophila* fly more erratically when density is low (Combes et al., 2012), suggesting that group dynamics affect flight behavior. From our data it is therefore unclear what cues triggered pursuit start. Since all our experiments used a black bead moving against a brighter background (Fig. 1A, Movie S1-3), it would be interesting to determine if this dark contrast is an important driver.

When blowflies pursue a bead they sometimes follow it for a long time, during which they keep a fixed distance to the bead, by controlling their forward speed based on the target’s angular size (Boeddeker et al., 2003), so that smaller beads (physical sizes) are followed at a closer distance. However, we did not see a similar relationship between bead size and distance (Fig. 5A, D), nor a correlation between hoverfly flight speed and the target’s angular size (dotted data, Fig. S3A, B). In contrast to blowflies, *S. pipiens* hoverflies control their forward speed based on the distance to the target (Collett and Land, 1975a), as do houseflies (Wehrhahn et al., 1982). We found only a weak correlation between hoverfly flight speed and distance to the bead (Fig. S3A-C).

It is unlikely that the lack of correlations was caused by technical limitations, such as our relatively low recording rate of 120 frames/s. Indeed, behavioral delays during target pursuit are often much longer than the 8.3 ms temporal resolution provided in our set-up. For example, when filmed at 1000 frames/s, predator steering changes have delays of 28 ms in the robber fly *Holcocephala*, 18 ms in the killer fly *Coenosia*, and 47 ms in the dragonfly *Plathemis* (Fabian et al., 2018; Mischiati et al., 2015). Furthermore, *Lucilia* blowflies display behavioral delays between 10 and 32 ms, when recorded at 190 frames/s (Varennes et al., 2020), which is close to the temporal resolution we used.

Previous work suggested that *E. tenax* males pursue targets traveling at female flight speeds (Collett and Land, 1978). However, in the field *Eristalis sp*. males pursue artificial targets moving at 5 – 12.5 m/s (Collett and Land, 1978), which is faster than typical female *Eristalis sp*. flight speeds (Thyselius et al., 2018). We here showed that hoverflies also pursue beads moving much slower than this, and even stationary targets (Fig. 3C). This is important since male *E. tenax* often wait for females to land before trying to mate with them (Fitzpatrick, 1981). Male *E. tenax* are capable of flying very fast, up to 10 m/s (Collett and Land, 1978). Indeed, even in our limited physical space, we found pursuit speeds at individual time points of up to 3.3 m/s (Fig. 5J). This could suggest that the males perceived fast moving beads as a male competitor rather than a cruising female. Furthermore, escaping female *E. tenax* can fly at up to 1.5 m/s (Thyselius et al., 2018), so the faster beads might have been perceived as escaping females.

### Pursuit style

Dragonfly and robber fly eyes often boast areas with improved spatial and temporal resolution, so called acute zones. They attempt to keep the target image in this acute zone during pursuit (Olberg et al., 2007; Wardill et al., 2017), a strategy shared with non-predatory dipterans, such as houseflies (Wagner, 1986; Wehrhahn et al., 1982) and the hoverfly *Syritta pipiens* (Collett and Land, 1975a). We found that male *E. tenax* pursue targets from above as well as from below (Fig. 4, 7). Male *E. tenax* harbor a dorso-frontal bright zone (Straw et al., 2006). Even if we did not reconstruct the head movements, the target image is unlikely to fall within the dorsal visual field when the hoverfly is flying above the bead (Fig. 4, 7E). However, the hoverflies attempted to keep the bead anterior relative to the flight direction, especially at the start of the proximal stage (Fig. 7D*iv*, E*iv*). The anterior visual field boasts higher resolution than the lateral visual field (Straw et al., 2006).

We broke down each pursuit into two stages, where the distal stage appeared to be optimized to rapidly decrease the distance to the target, and the proximal stage to staying close to the target (Fig. 5). Indeed, the distal stage could use proportional navigation based on the bearing angle, whereas the proximal stage did not (Fig. 7I-K). For *E. tenax* males the goal may not be to catch a target (Fig. 6), but to either chase it out of its territory if it is an intruder, or to stay close until it lands if it is a potential mate. Similar shadowing behavior has been described in dragonflies, previously referred to as motion camouflage (Mizutani et al., 2003). Indeed, staying close to the target allows the hoverfly to gather more information. In the field, *Eristalis sp*. males often chase intruders out of their territories without contact (Fitzpatrick, 1981). The males also often follow females, waiting for them to settle before mating, rather than grasping them in the air (Fitzpatrick, 1981). Indeed, we found that *E. tenax* males followed the artificial target for up to 3 s (Fig. 5H), and that when proximal the hoverfly’s translational speed was correlated with the bead speed (Fig. 5K), likely to stay in close vicinity, even if it was well within its capacity to speed up (see e.g. Fig. 5I) and catch the target. Indeed, they rarely got close enough to the 8 or 10 mm beads to suggest physical contact (Fig. 6). It might be beneficial for hoverflies to keep a longer distance to the target to avoid physical and potentially lethal contact. Could it thus be that *E. tenax* have developed a strategy that will take them close to but rarely in contact with their targets?

## Acknowledgments

We thank Annika Olsén, Mats Thyselius, Moa Thyselius, AB Cederholms Lantbruk, Louise Gustafsson, current and past lab members for valuable feedback during the many stages of this work.

## Data availability

All data and analysis scripts are available via DataDryad: https://datadryad.org/stash/share/zCoFLQZLDwiTrE8Kinvurdz79Kc8drQhJigu_6EGBKg

