## Supplementary material for "Hoverfly (*Eristalis tenax*) pursuit of artificial targets"

*[1]Department of Medical Cell Biology, Uppsala University, 75123 Uppsala, Sweden; [2]Flinders Health and Medical Research Institute, Flinders University, GPO Box 2100, Adelaide SA 5001, Australia; [3]College of Science and Engineering, Flinders University, GPO Box 2100, Adelaide SA 5001, Australia; [4]Department of Ecology, Evolution and Behavior, University of Minnesota, Saint Paul, MN 55108, USA*

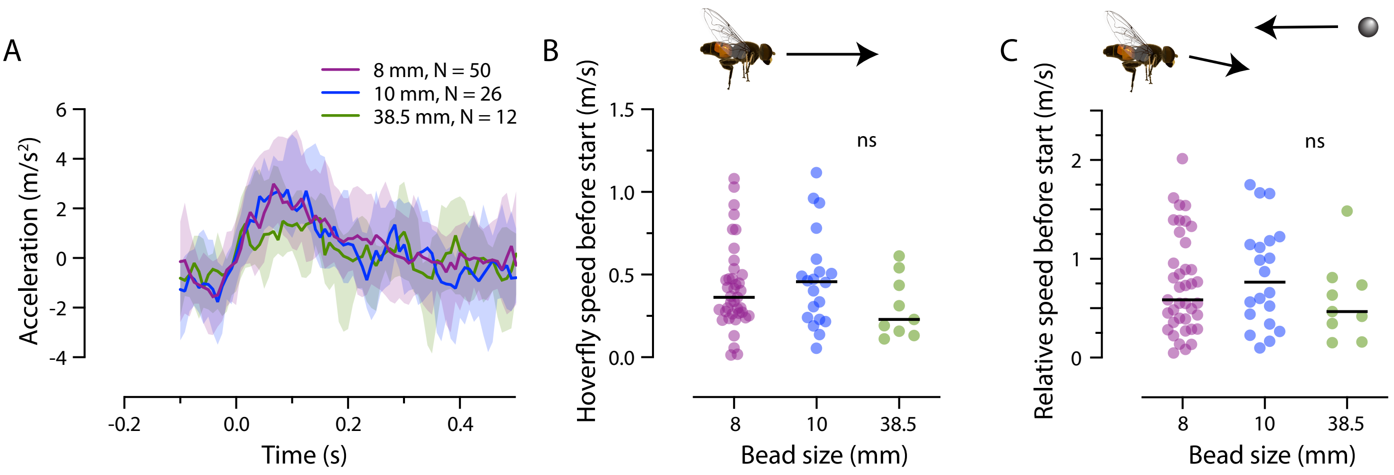


**Figure S1.** ***Hoverflies accelerate rapidly after pursuit start***
A) The hoverflies' acceleration at pursuit start. We smoothed the flight speed using a span of 10% of the total number of data points before calculating the acceleration. The data show the median +/- interquartile range, color coded according to bead size. There was no significant effect of bead size (using a mixed-effect model). B) The hoverfly flight speed 100 ms before pursuit start. C) The relative speed between the hoverfly and the bead 100 ms before pursuit start. The data in panels B and C were analyzed using Kruskal-Wallis tests.

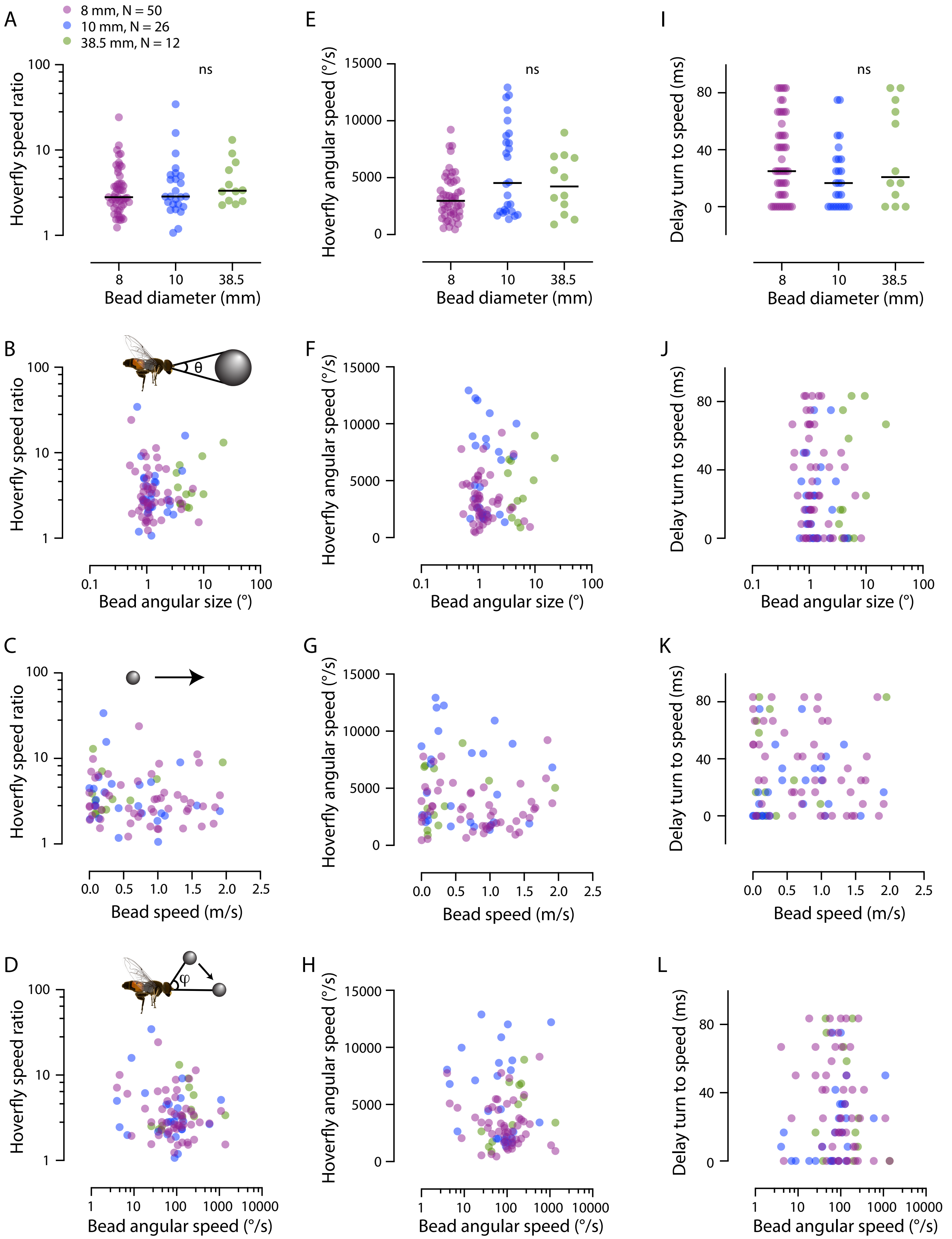

A
8 mm, N = 50
10 mm, N = 26
38.5 mm, N = 12
ns
Hoverfly speed ratio
Bead diameter (mm)
E
Hoverfly angular speed (°/s)
Bead diameter (mm)
I
Delay turn to speed (ms)
Bead diameter (mm)
B
θ
Bead angular size (°)
F
J
C
Bead speed (m/s)
G
K
D
φ
Bead angular speed (°/s)
H
L

***Figure S2. Hoverfly behavior at pursuit start does not depend on the bead***

Figure S2 aims to investigate how the hoverfly might adjust its flight behavior at pursuit start based on the bead. For this purpose, we display the hoverfly speed ratio (left column), the hoverfly angular speed (middle column), and the delay between the turn and speed increase (right column), as a function of the bead's physical size (top row), the bead's angular size (2$^{nd}$ row), the bead's translational speed (3$^{rd}$ row), and the bead's angular speed (bottom row). As evident from the data, there is no clear correlation between the hoverfly's behavior at pursuit start and any input from the bead. As explained in the main text, this is contrary to published accounts of pursuit behavior in the field (e.g. Collett and Land, 1978). The panels show the following: A) The hoverfly speed ratio following pursuit start, as a function of the bead diameter. B) The hoverfly speed ratio following pursuit start, as a function of the size of the bead's angular image. C) The hoverfly speed ratio, as a function of the bead's translational speed. D) The hoverfly speed ratio, as a function of the bead's angular speed. E) The hoverfly angular speed at pursuit start, as a function of bead diameter. F) The hoverfly angular speed at pursuit start, as a function of the bead's angular size. G) The hoverfly angular speed as a function of bead's speed at pursuit start. H) The hoverfly angular speed at pursuit start, as a function of the bead's angular speed. I) The delay between the hoverfly peak angular speed and peak speed ratio, as a function of bead diameter. J) The delay as a function of the bead's angular size. K) The delay as a function of the bead's speed. L) The delay as a function of the bead's angular speed. In all panels, the data are color coded according to bead size. The data in panels A, E, I were analyzed using Kruskal-Wallis tests.

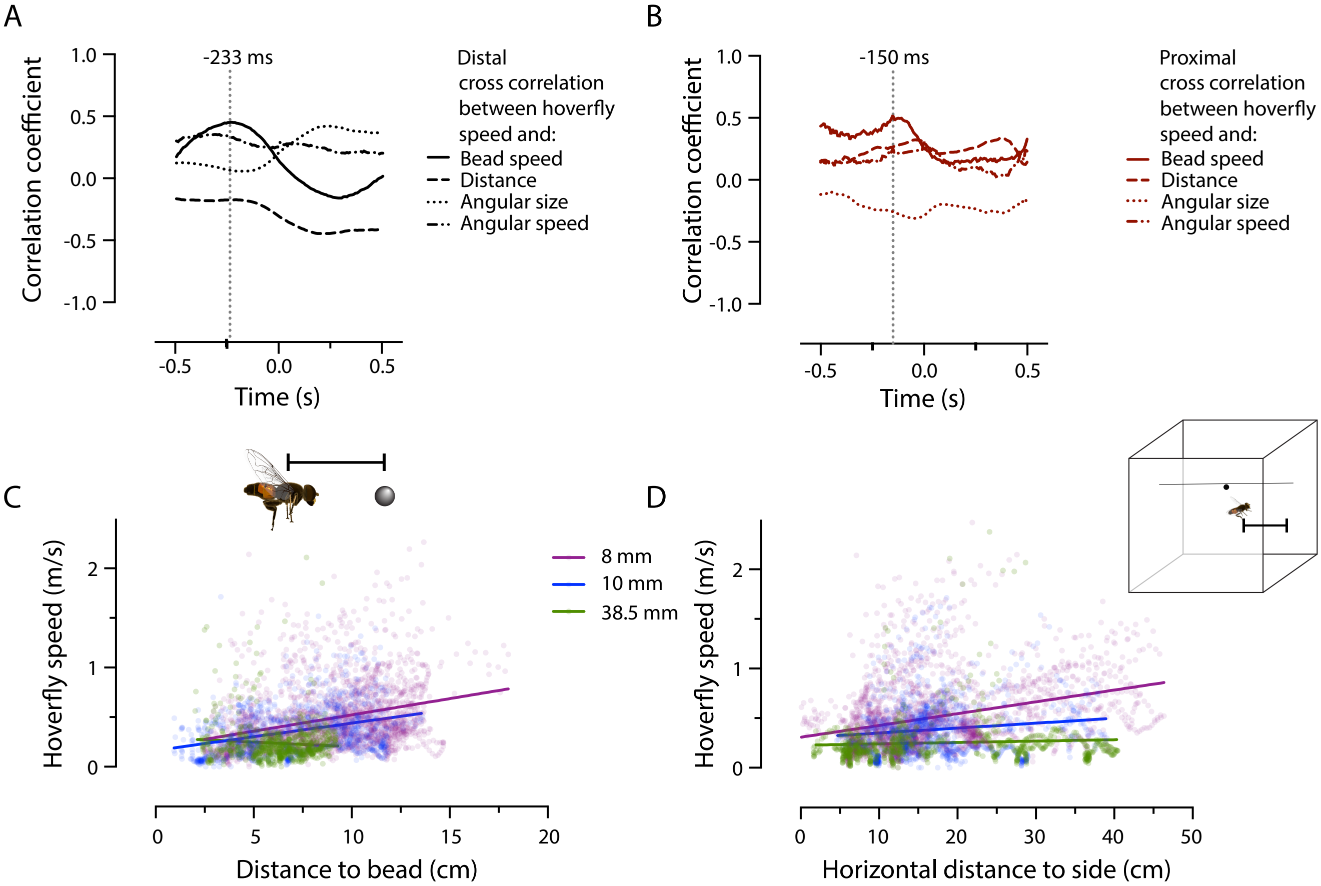


**Figure S3.** ***The hoverfly flight speed is best correlated with the bead speed***
A) The cross correlation between the hoverflies' translational speed and the beads' translational speed (solid line), the distance to the bead (dashed), the bead's angular size (dotted) and angular speed (dash-dotted) during the first 1 s of the distal stage of the pursuit. Peak correlation was found with the bead speed at -233 ms (dotted vertical line). B) Cross correlation between the hoverflies' translational speed and the beads' speed (solid line), the distance (dashed), the angular size (dotted), and the angular speed (dash-dotted), during the first 1 s of the proximal stage of the pursuit. Peak correlation was found with bead speed at 150 ms (dotted vertical line). C) The hoverflies' instantaneous flight speed as a function of the distance to the bead speed 58 ms previously (based on peak correlation, dashed line, panel B), during the first 1 s of the proximal stage. The lines show linear regression ($R^2$ = 0.096, 0.11 and 0.006 for the three bead sizes). D) The hoverflies' instantaneous flight speed as a function of the distance to either the left or the right wall of the arena, whichever was closest, during the first 1 s of the proximal stage. The lines show linear regression ($R^2$ = 0.10, 0.016 and 0.004 for the three bead sizes).

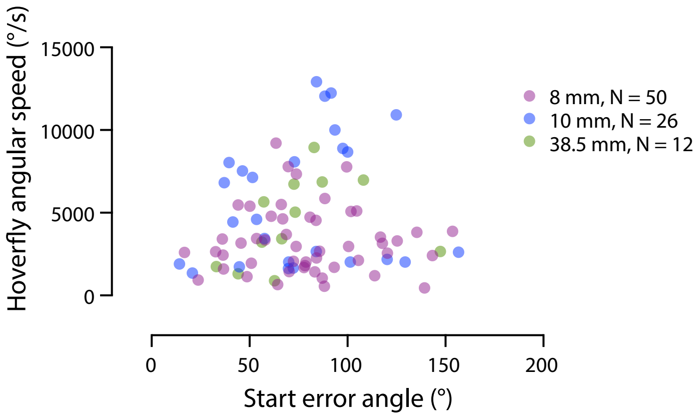


**Figure S4.** ***Hoverfly angular speed at pursuit start does not depend on the error angle to the bead***

The hoverfly angular speed, as a function of the error angle to the bead, both at pursuit start. The data are color coded according to bead size.

**Supplementary table**

**Table S1.** Hoverfly cohorts and their age when placed in the flight arena

| **Cohort #** | **Age of hoverflies at start** |
|---|---|
| 1 | 0-1 days |
| 2 | 5 months |
| 3 | 2-3 months |
| 4 | 3 weeks |
| 5 | 4 days |

**Supplementary movie legends**

***Movie S1. Pursuit of a 6 mm diameter bead***

The movie shows an example of a male *E. tenax* (pink circle) pursuing a 6 mm diameter bead (black circle). Pursuit start is highlighted with a yellow star.

***Movie S2. Pursuit of an 8 mm diameter bead, camera 1 view***

The movie shows an example of a male *E. tenax* (pink circle, which becomes purple during the proximal stage) pursuing an 8 mm diameter bead (black circle) programmed to move at 1 m/s. Pursuit start is highlighted with a yellow star. This pursuit is also shown in Figure 2B.

***Movie S3. Pursuit of an 8 mm diameter bead, camera 2 view***

The movie shows the same male *E. tenax* (pink circle, which becomes purple during the proximal stage) pursuing an 8 mm diameter bead (black circle) programmed to move at 1 m/s, seen from the second camera. Pursuit start is highlighted with a yellow star. This pursuit is also shown in Figure 2B.
